# Changes in the tumor microenvironment and treatment outcome in glioblastoma: A pilot study

**DOI:** 10.1101/2020.02.03.932475

**Authors:** Sehar Ali, Thaiz F Borin, Raziye Piranlioglu, Roxan Ara, Iryna Lebedyeva, Kartik Angara, Bhagelu R Achyut, Ali S. Arbab, Mohammad H Rashid

## Abstract

Glioblastoma (GBM) is a hypervascular and aggressive primary malignant tumor of the central nervous system. Recent investigations showed that traditional therapies along with antiangiogenic therapies failed due to the development of post-therapy resistant and recurrent GBM. Our investigations show that there are changes in the cellular and metabolic compositions in the tumor microenvironment (TME). It can be said that tumor cell-directed therapies are ineffective and we need to rethink how to treat GBM.

We hypothesize that the composition of TME-associated cells will be different based on the therapy and therapeutic agents, and TME-targeting therapy will be better to decrease recurrence and improve survival. Therefore, the purpose of this study is to determine the changes in the TME in respect of T-cell population, M1 and M2 macrophage polarization status, and MDSC population following different treatments in a syngeneic model of GBM. In addition to these parameters, tumor growth and survival were also studied following different treatments.

The results showed that changes in the TME-associated cells were dependent on the therapeutic agents and the TME-targeting therapy improved the survival of the GBM bearing animals.

The current GBM therapies should be revisited to add agents to prevent the accumulation of bone marrow-derived cells in the TME or to prevent the effect of immune-suppressive myeloid cells in causing alternative neovascularization, the revival of glioma stem cells, and recurrence. Instead of concurrent therapy, a sequential strategy would be best to target TME-associated cells.

## Introduction

Even with current treatment strategies and the addition of expensive immunotherapies or antiangiogenic therapies, the prognosis of glioblastoma (GBM) is dismal (1-3). GBM is a very hypervascular and invasive malignant tumor. So much so that, current treatments consisting of surgery, radiation and chemotherapies with or without adjuvant still show no hope to patients (4-6). Interestingly, recent investigations demonstrated that traditional therapies along with newer antiangiogenic therapies are changing the cellular as well as the metabolic compositions of the tumor microenvironment (**TME**) tremendously (7-11). Therefore, newer treatment strategies targeting TME should be considered along with targeting tumor cells in GBM.

The TME is composed of tumor cells, stromal cells, cells from the bone marrow, and the extracellular matrix (12). Except for a few cell types, such as normal epithelial cells, myoepithelial cells, dendritic cells, M1 macrophages, N1 neutrophils and CD8 T-cells, most of the stromal and bone marrow-derived cells promote tumor growth and metastasis (10, 11, 13-15). In fact, platelets have also been shown to promote tumor growth (16-19). Therefore, it is imperative to include targeting tumor-associated cells in the current standard regimen of therapies for malignant tumors such as GBM. However, there have been limited investigations done to understand the changes in the TME following standard as well as experimental therapies in GBM.

Tumor induction and evolution is driven by the interplay between stromal and immune cells within the TME. Tumor-associated macrophages (TAM), a critical component of the TME, have a differential function in respect to tumor growth and metastasis (20-22). TAM recruitment, localization, and phenotypes are regulated by the tumor-secreted factors at the hypoxic areas of the tumor (23, 24). Depending on the stimuli, macrophages undergo a series of functional reprogramming as described by two different polarization states, known as M1 and M2 (24, 25). Phenotypically, M1 macrophages express high levels of major histocompatibility complex class II (MHC II), the CD68 marker, and co-stimulatory molecules CD80 and CD86. On the other hand, M2 macrophages express high levels of MHC II, CD163, CD206/MRC1, Arg-1 (mouse only) and others. In the TME, classically activated macrophages, also known as M1 macrophages, are activated by tumor-derived cytokines such as granulocyte monocyte colony stimulating factor (GM-CSF), interferon-γ, and tumor necrosis factor (TNF). These M1 macrophages play an important role as inducer and effector cells in polarized type 1 helper T cell (Th1) responses. These Th1 cells drive cellular immunity to eliminate cancerous cells. To accomplish Th1 activation, M1 macrophages produce high amounts of IL-12 and IL-23, and low amounts of IL-10, reactive oxygen and nitrogen species, and IL-1β, TNF, and IL-6 inflammatory cytokines (25, 26). M1 macrophages also release anti-tumor chemokines and chemokines such as CXCL-9 and CXCL-10 that attract Th1 cells, (27-29). Th1 cells drive cellular immunity to eliminate cancerous cells. On the other hand, M2-polarized macrophages, also known as alternatively activated macrophages are induced by IL-4, IL-13, IL-21 and IL-33 cytokines in the TME (30, 31). M2 macrophages release high levels of IL-10 and, transforming growth factor-beta (TGF-β) and low levels of IL-12 and IL-23 (type 2 cytokines). M2 macrophages also produce CCL-17, CCL-22, and CCL-24 chemokines that regulate the recruitment of Tregs, Th2, eosinophils, and basophils (type-2 pathway) in tumors (27, 29). The Th2 response is associated with the anti-inflammatory and immunosuppressive microenvironment, which promotes tumor growth.

Recent investigations including our own indicated the involvement of myeloid-derived suppressor cells (MDSCs) in the primary as well as metastatic TME (32-36). MDSCs are a heterogeneous population of immature myeloid cells, generated from bone marrow hematopoietic precursor cells that fail to undergo terminal differentiation to mature monocytes or granulocytes. They are divided broadly into monocytic (CD11b+/Gr1+/Ly6C+) and granulocytic (CD11b+/Gr1+/Ly6G+) (37-39). During tumor progression, MDSCs are greatly expanded and they exhibit remarkable immunosuppressive and tumorigenic activities. They are directly implicated in the escalation of tumor metastases by partaking in the epithelial-mesenchymal transition (EMT) and, tumor cell invasion, while also promoting angiogenesis and formation of the pre-metastatic niche (13, 33, 34). MDSCs were demonstrated to promote tumor invasion and metastasis by two mechanisms: (i) increasing production of multiple matrix metalloproteinases (MMPs) that degrade the extra-cellular matrix and chemokines that establish a pre-metastatic milieu (40, 41), and (ii) merging with tumor cells (42, 43).

From the above discussion, it is obvious that TME-associated bone marrow-derived cells are important in treatment resistance, invasion and metastasis. Therefore, the purpose of this study is to determine the changes in the TME in respect of T-cell population, M1 and M2 macrophage polarization status, and MDSC population following different treatments in a syngeneic model of GBM. In addition to these parameters, tumor growth and survival were also studied following different treatments. In this study, we have used the following agents: a drug that alters hydroxylase pathways of arachidonic acid metabolism (HET0016 and its different analogs), colony stimulating factor 1 receptor (CSF1R) inhibitor (GW2580), anti PD-1 (program death) antibody, CXCR2 receptor blockers (Navarixin and SB225002), temozolomide (TMZ), irradiation, VEGFR2 receptor tyrosine kinase inhibitor (Vatalanib), and conditional CSF1R knockout mice plus different treatments.

## Materials and methods

### Ethics statement

All the experiments were performed according to the National Institutes of Health (NIH) guidelines and regulations. The Institutional Animal Care and Use Committee (IACUC) of Augusta University (protocol #2014–0625) approved all the experimental procedures. All animals were kept under regular barrier conditions at room temperature with exposure to light for 12 hours and dark for 12 hours. Food and water were offered ad libitum. All efforts were made to ameliorate the suffering of animals. CO2 with a secondary method was used to euthanize animals for tissue collection.

### Materials

HPßCD (2-hydroxy Propyl-β-Cyclodextrin) was purchased from Sigma-Aldrich (St. Louis, MO), cell culture media was from Thermo Scientific (Waltham, MA), and fetal bovine serum was purchased from Hyclone (Logan, Utah). HET0016 was made by Dr. Levedyeva in the Department of Chemistry, Augusta University with a purity of more than 97%. Cell culture grade DMSO was purchased from Fischer Scientific (PA). We made the complex of HET0016 plus HPßCD as per our previous publication (8). VEGFR2 tyrosine kinase inhibitor (Vatalanib) and colony stimulating factor 1 receptor (CSF1R) inhibitor (GW2580) were purchased from LC Laboratories, Woburn, MA. SB225002 (CXCR2 inhibitor) was purchased from Selleckchem, Houston, TX. Navarixin was purchased from MedKoo bioscience Inc, Morrisville, NC. All flow antibodies are from Bio Legend, San Diego, CA. All antibodies for western blotting, immunohistochemistry, and immunofluorescence were purchased from Santa Cruz (total-CXCR2 and anti-GAPDH), R&D systems (anti-hCXCR2), Thermo Scientific (anti-Laminin), and Sigma Aldrich (β-actin and FITC-conjugated tomato lectin). All culture media were purchased from Corning and GE Healthcare Life Sciences.

### Tumor cells and orthotopic animal model of GBM

To determine the *in vivo* effect of different treatments, orthotopic GBM models using syngeneic GL261 cells in wild type and CSF1R conditional knockout C57BL/6 mice were prepared according to our published methods (8, 10, 11, 44). In short, luciferase positive GL261 cells were grown in standard growth media (RPMI-1640 plus 10% FBS) and collected in serum-free media on the day of implantation. After preparation and drilling a hole at 2.25 mm (athymic nude mice) to the right and 2 mm posterior to the bregma, taking care not to penetrate the dura, a 10 µL Hamilton syringe with a 26G-needle containing tumor cells (10,000) in a volume of 3 µl was lowered to a depth of 4 mm and then raised to a depth of 3 mm. During and after the injection, a careful note was made for any reflux from the injection site. After completing the injection, we waited 2-3 minutes before withdrawing the needle 1 mm at a time in a stepwise manner. The surgical hole was sealed with bone wax. Finally, the skull was swabbed with betadine before suturing the skin (45-47). There were at least three animals in each group of treatment. Tumor growth was determined by optical imaging (bioluminescence imaging after injecting luciferin) on days 8, 15 and 22. For flow cytometry of tumor-associated cells, animals were euthanized on day 22 after the last optical imaging. Both male and female animals were used.

### Treatments

All treatments were started on day 8 following tumor implantation and continued for two weeks. The following treatment groups were used to determine the TME associated T-cells, different macrophages, MDSCs present by flow cytometry; 1) vehicle, 2) HET0016 complexed with HPßCD at 10mg/kg/day for 5 days/week, intravenous (IV), 3) GW2580, 160mg/kg/day 3day/week, oral, 4) temozolomide (TMZ) 50mg/kg/day, 3days/week, oral, 5) Vatalanib 50mg/kg/day, 5 days/week, oral, 6) Navarixin, 10mg/kg/day, 5 days/week, intraperitoneal (IP), 7) anti-PD-1 antibody, 200µg/dose, 2 doses/week, IP, 8) image guided radiation therapy, 10Gy/dose/week for two weeks, 9) combined HET0016 plus GW2580, 10) combined HET0016 plus GW2580 plus anti-PD-1 antibody.

### Making of a conditional knockout mouse model of bone marrow-derived CSF1R+ myeloid cells

Heterozygous CSF1R _flox/wt_/MX1-Cre+ male was mated with a heterozygous CSF1R _flox/wt_/MX1-Cre+ female to achieve 25% of the progeny with homozygous CSF1R_flox/flox_/MX1-Cre+ (knockout) genotype in bone marrow cells. Other progeny was wild-type CSF1R_wt/wt_/MX-1-Cre+ (25%) and heterozygous CSF1R_flox/wt_/MX-1-Cre+ (50%) genotypes. After repeated cross-breeding, we have generated a colony of CSF1R_flox/flox_/Cre+ (knockout). These animals are healthy and are being used for breeding. Analysis of myeloid cells in the peripheral blood before and after injection of polyinosinic-polycytidylic acid (poly-IC) showed bone marrow-specific depletion of CSF1R+ cells (**Figure 1**). These animals (male and female) were used to generate GL261 derived syngeneic GBM after depletion of bone marrow-derived myeloid cells and then treated with HET0016 or anti PD-1 antibody alone or in combination or with CXCR2 antagonist SB225002 (10mg/kg/day 5 days/week, IP) for two weeks.

**Figure 1:**
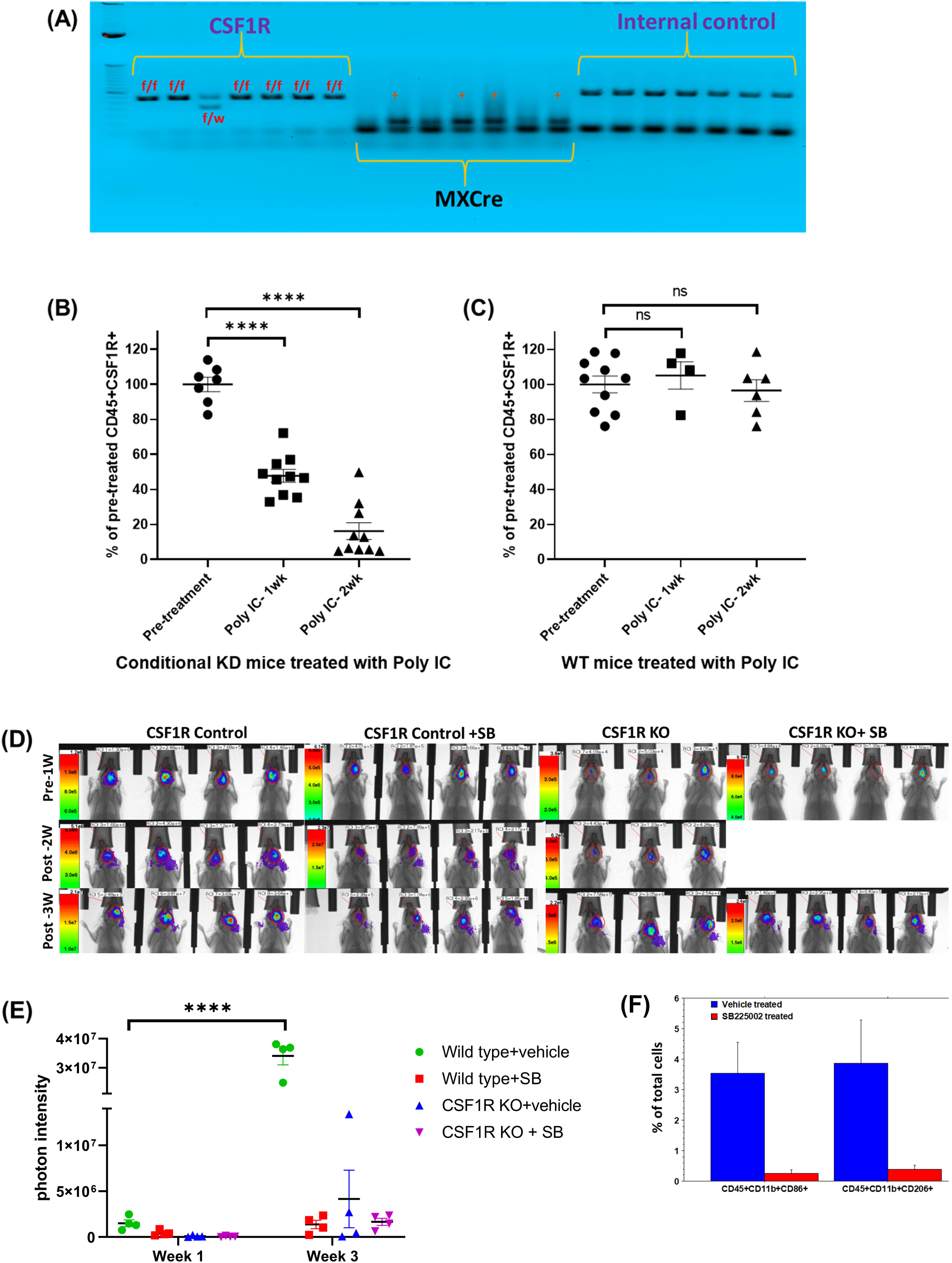
CSF1R conditional knockout mouse and GBM development. **(A)** Agarose gel electrophoresis showing homozygous CSF1R^flox/flox^/MX1-Cre+ (knockout) genotype. **(B)** Flow-cytometric analysis of peripheral blood cells from conditional knock out mice showed a significant dose-dependent decrease in CD45+CSF1R+ cells following two weeks of treatments with poly-IC. **(C)** Flow-cytometric analysis of peripheral blood cells from wild type mice did not show any significant difference in CD45+CSF1R+ cells following two weeks of treatments with poly-IC. **(D and E)** Optical images and quantified photon intensities of pre and post-treatment (either vehicle or SB225002) showed significantly increased tumor growth in the vehicle-treated wild type animals after 3 weeks. Knock out animals treated with either vehicle or SB225002 and wild type animals treated with SB225002 did not show any significant tumor growth after 3 weeks. **(F)** Flow-cytometric analysis showing significantly decreased tumor-associated CD45+CD11b+CD86+ and CD45+CD11b+CD206+ cells.

### Determination of bone marrow-derived cells in the TME

Following euthanasia, animals were perfused with ice-cold PBS and the right brain containing GBM was collected, passed through 40-micron mesh and a single-cell suspension was made. Similarly, spleens were collected, passed through 40micron mesh and a single-cell suspension was made. Before adding panels of antibody cocktail, non-specific uptake of the antibody was blocked by adding recommended blocker. The population of the following cells were determined by a Accuri C6 flow cytometer from cells collected from tumors and spleen; CD45+/CD4+, CD45+/CD8+, CD45+/CD11B+/Gr1+/Ly6C+, CD45+/CD11B+/Gr1+/Ly6G+, CD45+/CD86+/CD80+, AND CD45+/CD206+. The findings were compared among all the treatment groups.

### Determination of tumor growth

Bioluminescent imaging was used to determine the tumor growth following different treatments. All animals underwent imaging following IP injection of luciferin (150mg/kg). Images were obtained from all animals on days 8, 15 and 22. Photon density (photon/sec/mm^2^) was determined by drawing an irregular region of interest to cover the tumor area. The findings were compared among all the treatment groups.

### Determination of survival

Groups of animals were also used to determine the survival following different TME targeted therapies. All animals were routinely observed 2-3 times a week to assess the wellbeing as well as body weight. The animals were followed up until they become moribund or fulfill the criteria for euthanasia as per the approved IACUC protocols. The findings were compared among all the treatment groups.

### Statistical analysis

Quantitative data were expressed as mean ± standard error of the mean (SEM) unless otherwise stated. For the flow-cytometric studies, we used ordinary one-way analysis of variance (ANOVA) followed by multiple comparisons using Dunnett’s multiple comparisons test. For BLI (optical imaging) data, the general framework of analyses included two-way ANOVA followed by either Tukey’s or Sidak’s multiple comparisons. We analyzed the survival of the animals following different treatments. Log-rank test (Mantel-Cox) was applied to determine the significance of differences among the groups. A P value of 0.05 was considered significant.

## Results

In this study, we successfully developed CSF1R conditional knockout mouse. These conditional knock out mice showed homozygous CSF1R^flox/flox^ /MX1-Cre+ (knockout) genotype (**Figure 1A**). Compared to wild type mice, conditional knockout mice showed a significant dose-dependent decrease in CD45+CSF1R+ cells following two weeks of treatments with poly-IC. There was almost 80% decrease of CSF1R+ cells in the peripheral blood (**Figure 1B**). Wild type mice treated with poly-IC did not show any significant difference in CD45+CSF1R+ cells (**Figure 1C**). Both wild type (control) and knockout mice (after two weeks’ of treatments with poly-IC) received intracranial implantation of syngeneic GL261 glioblastoma. On day 8 of tumor implantation, groups of animals received either vehicle or SB225002 for two weeks. All animals underwent optical imaging pre and post-treatment. Photon intensities were determined to measure tumor growth. Wild type control animals showed significantly increased tumor growth (**Figure 1D**) which is indicated by a 10-fold increase in the photon intensity (**Figure 1E**). On the other hand, both wild type (control) treated with SB225002 and knockout mice showed significantly decreased tumor growth at week 3, indicating the involvement of CSF1R+ cells in the TME. It is also known that the CXCR2 antagonist can inhibit the function of myeloid cells by blocking the interaction of CXCR2 and IL-8 (48-50). Tumor-associated CD45+CD11b+CD86+ and CD45+CD11b+CD206+ cells were determined following treatment with SB225002 in wild type animals. Both cell types were significantly decreased following the treatments (**Figure 1F**). T-cells and MDSC populations showed no significant difference between the treated and untreated wild type animals.

Both wild type and CSF1R knockout mice received different treatments that target tumor cells or tumor-associated cells. All treatments were for two weeks and the treatment was started on day 8 of orthotopic tumor implantation. On day 22 following last optical imaging, animals were euthanized and the tumors were collected for flow cytometry to determine the population of T-cells (CD4, CD8), CD11b+ cells, macrophages (M1 and M2), and MDSCs (Ly6C and Ly6G). To our surprise, CD4, CD8, CD11b, and Ly6G positive cells significantly increased in tumors treated with TMZ (**Figure 2**). On the other hand, different cellular populations were significantly decreased in post-radiation tumors. All other treatments that targeted tumor-associated myeloid cells or checkpoint showed increased accumulation of CD4 and CD8 cells in the tumors but myeloid cell populations including MDSCs, CD11b+ cells, and macrophages showed insignificant changes in the TME compared to that of control and Vatalanib treated tumors (**Figure 3**).

**Figure 2:**
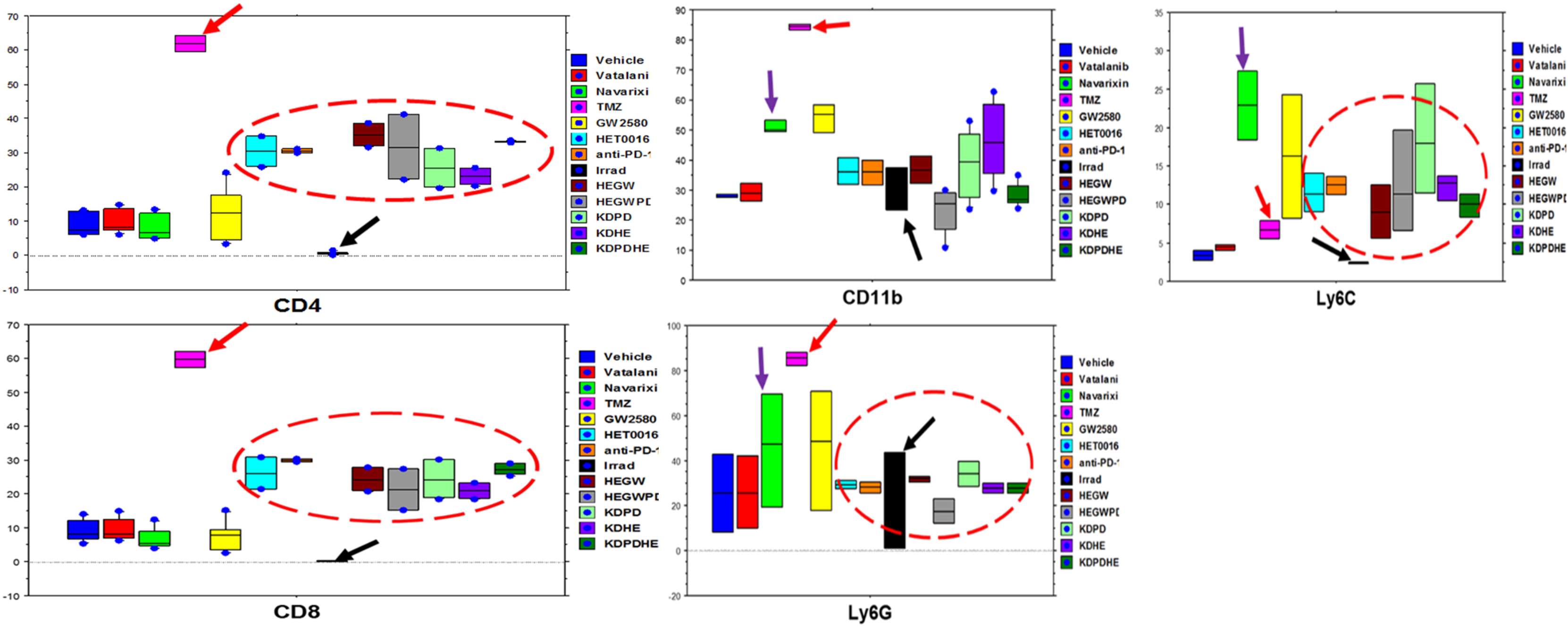
Flow cytometric analysis of T-cells and myeloid cell populations in wild type and knockout animals. There was a significant increase in CD4, CD8, CD11b, and Ly6G positive cells in tumors treated with TMZ (red arrows) while irradiation caused a significant reduction (black arrows) in different cellular populations compared to control group. All other treatments showed increased infiltration of CD4 and CD8 T-cells but insignificant changes in MDSCs, CD11b populations.

**Figure 3:**
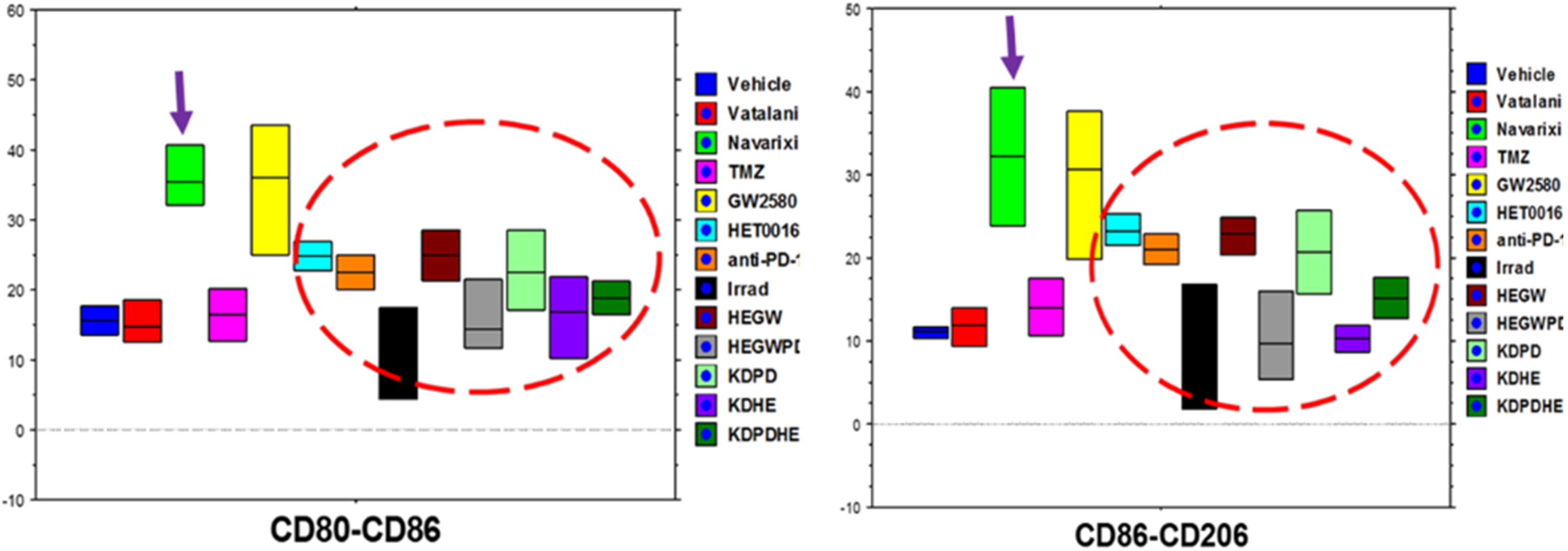
Flowcytometric analysis of M1 and M2 macrophage populations. Treatment with Navarixin and GW2580 increased the macrophage population insignificantly, and all other treatments changed the macrophage population inconsequentially.

All animals that were followed for survival and euthanized on day 22 to determine the TME associated cells also underwent optical imaging before treatment and at one and two weeks after treatments. The dose of luciferin and exposure time were kept identical for every animal at each time point. Then the photon intensity (intensity/sec/mm^2^) was determined by making an irregular region of interest encircling the tumors at each time point. **Figure 4** shows the tumor growth following different treatments. Tumor in all therapy groups except in Vatalanib treated animals, were stable following 1 week of treatments and there was no significant difference compared to that of vehicle-treated animals. However, Vatalanib treated animals showed significantly increased photon intensity indicating tumor growth following 1 week of treatments. Tumor growths were substantially increased in vehicle, Vatalanib, and TMZ treated animals following 2 weeks of therapy indicating the development of resistance in TMZ group. All other groups showed increased tumor growth but were significantly slower than that of vehicle, Vatalanib, or TMZ treated animals. It should be noted that the animals that received TME-associated cell-directed therapy showed significantly lower tumor growth 2 weeks following treatments. The animals that receive antiangiogenic (Vatalanib) and tumor cell-targeted (TMZ) therapy exhibited rebound tumor growth at 2 weeks of treatments.

**Figure 4:**
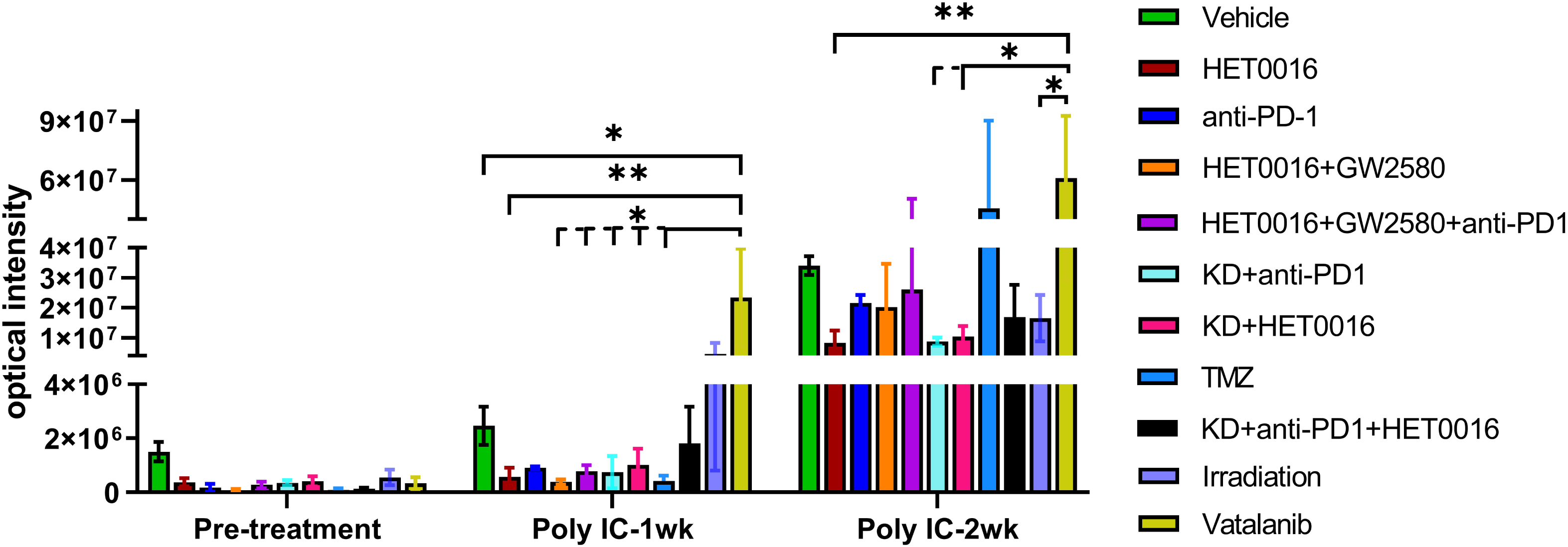
Bioluminescent image-based analysis of tumor growth. All animals underwent optical imaging to monitor tumor growth before starting the treatment (day 8 post-inoculation), 1 week, and 2 weeks after treatment. There was no significant difference between all treatment groups compared to that of vehicle-treated animals after 1 week of treatment except the Vatanallib treated group that showed significant tumor growth. Following 2 weeks of treatment, tumor growths were substantially increased in the vehicle, Vatalanib, and TMZ treated animals. All other groups showed increased tumor growth but were significantly slower than the above-mentioned groups.

We instituted different treatments targeting both tumor cells and the tumor microenvironment including arachidonic acid metabolisms and anti-depressant (selective serotonin reuptake inhibitor (SSRI), fluoxetine) drugs alone or in combination with TMZ. We also used a very high dose of HET0016 (50mg/kg/day). Usual dose of HET0016 is 10mg/kg/day. All treatments significantly increased the survival of animals bearing syngeneic GL261 GBM (**Figure 5A**). The most significantly increased survival was observed in animals’ groups that were treated with TMZ, HET0016, TMZ+HET0016, and with a HET analog. Although Navarixin (IL-8CXCR2 axis blocker) increased the survival of the animals, the addition of TMZ did not improve survival (**Figure 5B**).

**Figure 5:**
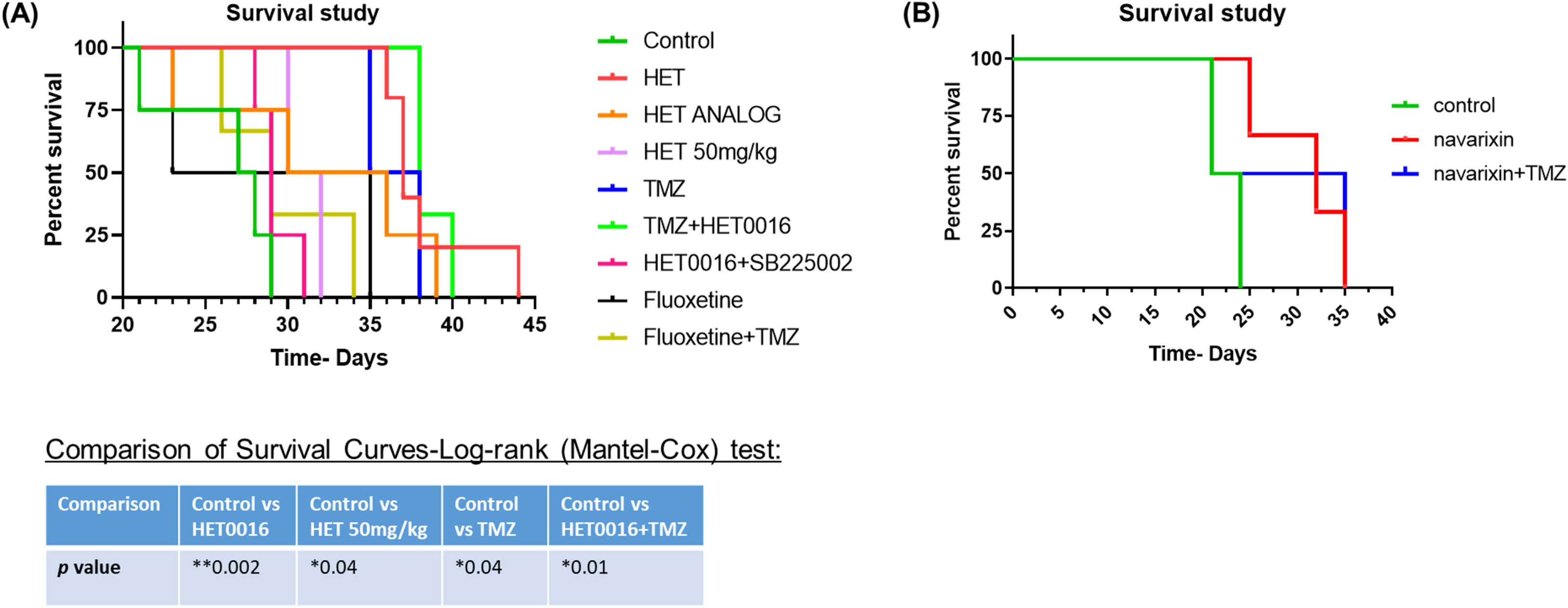
Survival studies showing improved survival following the use of TME targeting agents. **(A and B)** Kaplan-Meier curve showing significantly increased survival in animal groups treated with TMZ, HET0016, TMZ+HET0016, and with a HET analog. Although Navarixin) increased survival, the addition of TMZ with it did not improve the outcome.

## Discussion

GBM is a devastating malignant tumor of the central nervous system. Once diagnosed it becomes a death sentence to patients within 15 months (51-54). Currently, surgical resection followed by radiation and TMZ therapies is the standard of care for GBM patients (55). With these extensive therapies, almost all patients show therapy resistance and recurrence of GBM (56). To address resistance and recurrence, clinicians have adopted antiangiogenic therapies in recurrent GBM. These treatments decrease the formation of new blood vessels and decrease edema, thus reducing the dose of corticosteroids needed after therapy (57, 58). Additionally, advanced immunotherapy and targeted therapies have been instituted (59). However, early reports demonstrated that these are non-effective treatment strategies (10, 45, 60-65). Investigations from our lab indicated that most of the instituted therapies mobilized bone-marrow cells to the sites of GBM and orchestrated therapy resistance (10, 11). Our results showed that antiangiogenic therapies initiate alternate vascularization pathways and eventually increased neovascularization in therapy-resistant GBM (7, 45, 66). We found that angiogenic and vasculogenic myeloid cells accumulated at GBM sites following therapies(11, 65). Furthermore, we reported the process of vascular mimicry in which GBM cells transdifferentiate into glioma stem cells that can then form functional blood vessels (7, 67). All of these results support our conclusion that the possible changes occurring in the TME following standard or investigational treatments in GBM have not been properly studied. This includes both changes in TME associated cells as well as the changes that occur in the metabolic cascade of TME associated cells. In this pilot study, we aimed to investigate these changes. To accomplish this, we used standard therapies (radiation and TMZ) as well as agents that targeted TME associated cells (CSF1R inhibitor GW2580 to target myeloid cells, IL-8-CXCR2 antagonists Navarixin and SB225002 to target stem cells causing vascular mimicry, anti-PD1 antibody targeting immune suppressive molecules) and different metabolic pathways (HET0016 and its analog to target CYP4A-20-HETE axis of arachidonic acid metabolisms, fluoxetine to target serotonin reuptake). Following therapies, we determined the changes in the composition of TME-associated cells and the survival benefit of the therapeutic agents alone or in combination with TMZ.

Our results clearly demonstrated the importance of TME associated CSF1R positive cells. Animals treated with GW2580 and conditional knockout animals (CSF1R knockout) showed a decreased number of myeloid cells in the TME, whereas TMZ therapy increased the population of myeloid cells in the treated GBM. Previously, our reported results, as well as results from different investigators, have proven the importance of myeloid cells in developing therapy resistance in GBM and other cancers (11, 13, 15, 68-70). Myeloid cells, such as macrophages and MDSCs, produce an immunosuppressive microenvironment that promotes tumor growth. Following chemotherapy, macrophage differentiation is altered to promote the production of cancer-supporting M2 macrophages in the TME (71). Chemotherapy has also been shown to promote macrophage aggregation, thus facilitating cathepsin protease B- and S-mediated therapy resistance (72). Some chemotherapeutic agents activate MDSCs to produce IL-1β. This leads to the secretion of IL-17 by CD4^+^ T-cells (73). Additionally, MDSCs have been shown to partake in the epithelial-mesenchymal transition, increase the production of multiple matrix metalloproteinases, and merge tumor cells (71-73). Therefore, the addition of myeloid cell blockage could mitigate these mechanisms of resistance. However, it is to note that, previous investigations also indicated the development of resistance following long-term therapy using CSF1R inhibitors (74, 75). This indicates the importance of sequential or intermittent therapy targeting GBM TME associated cells following or in between standard therapies for GBM.

To our surprise, we noticed a decreased accumulation of T-cells as well as different myeloid cell populations in the TME following radiation therapies. This decreased accumulation of T-cells may be due to the disruption of intact blood vessels that act as a delivery system of T-cells to the tumor site. This disruption is likely caused by radiation therapy-induced necrosis in tumors leading to tumor cell death. Therefore, most tumor recurrence in post-radiation GBM occurs from the periphery of the irradiated areas where a few cells may have survived the radiation injury. Our previous studies showed that the addition of HET0016 (blocker of CYP4A-20-HETE axis of arachidonic acid metabolisms) improved the survival of animals bearing patient-derived xenograft (PDX) GBM following 30Gy of radiotherapy (8). HET0016 is known to inhibit tumor and endothelial cell (EC) proliferation, EC migration, and prevent neovascularization including vascular mimicry (44, 67, 76). Although we have not tested agents that prevent the repair of DNA damage, the addition of PARP inhibitor may also help prevent the recurrence of GBM following radiotherapy (77, 78). However, in contrast to HET0016, PARP inhibitor has a very narrow therapeutic window and causes severe toxicity (77). Therefore, adding an inhibitor of arachidonic acid metabolic pathways may be useful in preventing the recurrence of post-radiation GBM.

Previously, we have reported the effectiveness of HET0016 in controlling GBM and breast cancer (8, 32). However, we had not yet reported TME-associated cells present following the treatment of HET0016. In this study, HET0016 treatment exhibited a similar phenomenon to that of myeloid cell-targeted therapies. It showed an increased T-cell population in the TME compared to that of vehicle and Vatalanib treated GBM. There was also a tendency to decrease immunosuppressive myeloid cell populations in the TME. Additionally, treatments using HET0016 and its analog showed significantly improved survival which corroborates with our previous reports (8). Our ongoing investigations show that the CYP4A-20-HETE pathway is active not only in tumor cells but also in TME associated myeloid cells (data not shown). Inhibition of 20-HETE increases the cytotoxic T-cells population in *in vitro* studies (manuscript under preparation). Details of HET0016 mediated therapies and its mechanisms are discussed in our previous reports (8). Therefore, we propose that the use of an inhibitor of the cytochrome P450 γ-hydroxylase pathway of arachidonic acid metabolisms may be used as an agent to target post-therapy GBM to prevent recurrence.

## In conclusion

current GBM therapies should be revisited to add agents to prevent the accumulation of bone marrow-derived cells in the TME or to prevent the effect of immune-suppressive myeloid cells in causing alternative neovascularization, the revival of glioma stem cells, and recurrence. Instead of concurrent therapy, a sequential strategy would be best to target TME associated cells.

## Acknowledgment

The authors like to acknowledge the help of the core facility of small animal imaging (CIFSA) for acquiring optical images.

## Author Contributions Statement

*<u>Sehar Ali</u>*: Analyze the flow cytometry and optical image data. She also helped writing the manuscript.

*<u>Thaiz F Borin</u>*: Tumor cell propagation, tumor implantation and acquisition of flow cytometry data. She edited the manuscript.

*<u>Raziye Piranlioglu</u>*: Tumor cell propagation, tumor implantation and acquisition of flow cytometry data.

*<u>Roxan Ara</u>*: Help acquiring optical images and analysis

Iryna Lebedyeva: Synthesize HET0016 and its analog. Helped editing the manuscript.

*<u>Kartik Angara</u>*: Initiated the experiments using CXCR2 antagonist and Vatalanib treatments

*<u>Bhagelu R Achyut</u>*: Helped making the transgenic CSF1R knockout animals and conducted initial optimization of Poly-IC injection and depletion of CSF1R+ cells. He also helped the interpretation of TME data.

*<u>Ali S. Arbab</u>*: Conceived the hypothesis, design the experiments and provided the funds. He edited the manuscript.

*<u>Mohammad H Rashid</u>*: Helped Sehar Ali to analyze the data, interpreted the results, maintaining CSF1R knockout animal colony, tumor implantation, treating animals, acquisition of flow cytometry data, preparing graphs and wrote the manuscript.

## Contribution to the Field Statement

Glioblastoma (GBM) is a devastating primary brain cancer. Current treatments that use surgery, chemotherapy and radiotherapy do not increase the survival of the patient. Almost all patients with GBM die with 15 months of diagnosis. GBM is also a tumor with many blood vessels, therefore, clinician started using anti-neovascular agents. However, recent reports indicated that all these treatments caused therapy resistance and enhance alternative neovascularization due to mobilization and accumulation of cells derived from patients’ bone marrow. These mobilized bone marrow cells accumulate in the GBM microenvironment and initiate an environment that is immunosuppressive and increase tumor cell invasion causing recurrent tumors. There is a movement of rethinking of therapy strategies in GBM. Investigators started using immunotherapy to change the microenvironment, however, early results are not encouraging. We hypothesize that agents that target GBM microenvironment should be included along with standard therapies either concurrently or sequentially. In this studies we showed the changes in GBM microenvironment following different therapies and showed the improvement of survival in mouse model following GBM microenvironment targeting therapies.

